## Supplementary Figures for "Divergent features of the coenzyme Q:cytochrome *c* oxidoreductase complex in *Toxoplasma gondii* parasites"

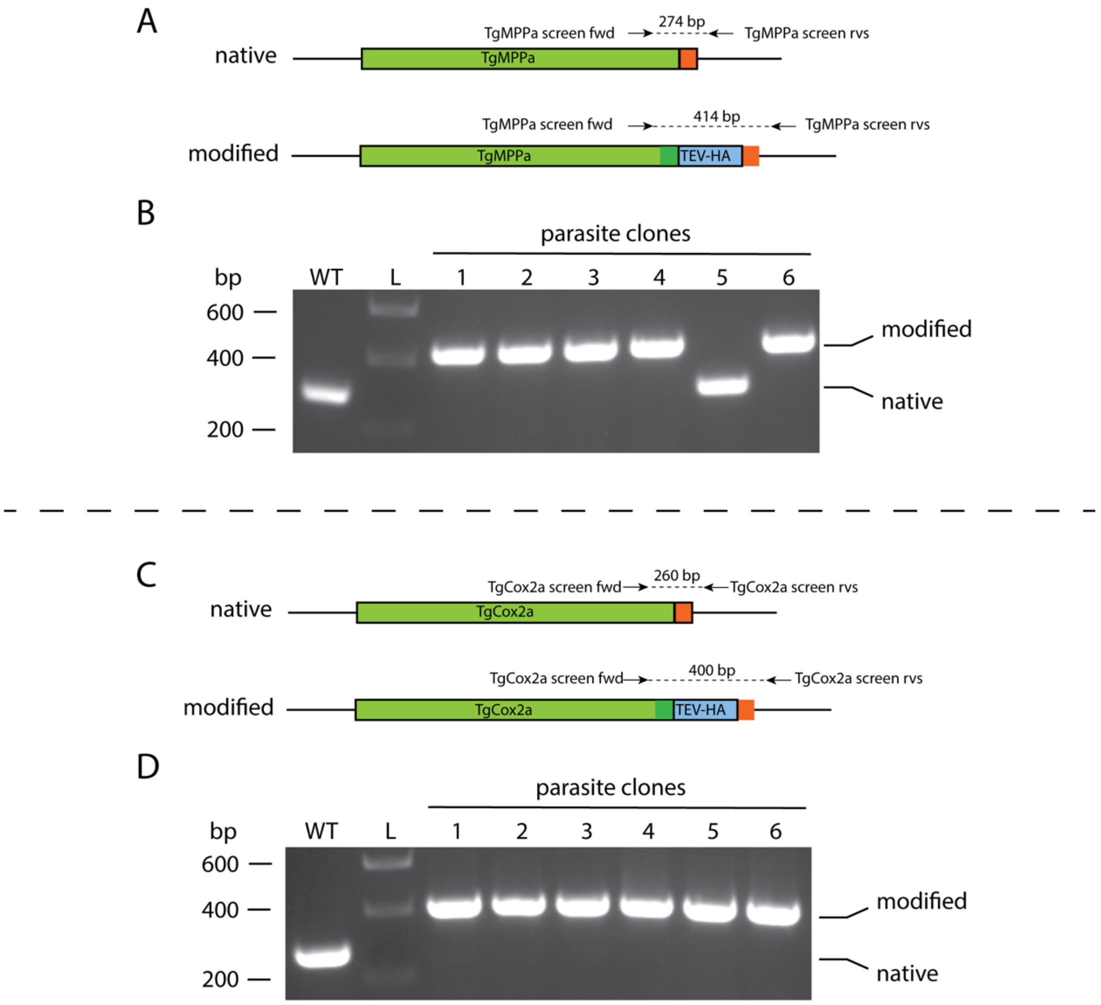


**Figure S1: Generating TEV-HA tagged *Tg*MPPα and *Tg*Cox2a strain parasites. (A)** Diagram depicting the 3' replacement strategy to generate TEV-HA-tagged *Tg*MPPα. A sgRNA was designed to target the *T. gondii* genome near the stop codon of *Tg*MPPα, and cause a double stranded break. A plasmid containing the sgRNA and GFP-tagged Cas9 endonuclease was co-transfected into *T. gondii* parasites with a PCR product encoding a TEV-HA epitope tag flanked by 50 bp of sequence homologous to the regions immediately up- and down-stream of the *Tg*MPPα stop codon. The homologous repair pathway of the parasite mediates integration of the PCR product into the *Tg*MPPα locus. Forward and reverse primers were used to screen parasite clones for successful integration of the TEV-HA tag at the target site, yielding a 274 bp product in the native locus and a 414 bp product in the modified locus. **(B)** PCR screening analysis using genomic DNA extracted from putative *Tg*MPPα-TEV-HA parasites (clones 1 – 6). Clones 1 – 4 and 6 yielded PCR products that indicated that these clones had been successfully modified. Genomic DNA extracted from wild type (WT) parasites was used as a control. **(C)** Diagram depicting the 3' replacement strategy to generate TEV-HA-tagged *Tg*Cox2a. A sgRNA was designed to target near the stop codon of *Tg*Cox2a. A plasmid containing the sgRNA and GFP-tagged Cas9 endonuclease was co-transfected into *T. gondii* parasites with a PCR product encoding a TEV-HA epitope tag flanked by 50 bp of sequence homologous to the regions immediately up- and down-stream of the *Tg*Cox2a stop codon. Forward and reverse primers were used to screen parasite clones for successful integration of the TEV-HA tag at the target site, yielding a 260 bp product in the native locus and a 400 bp product in the modified locus. **(D)** PCR screening analysis using genomic DNA extracted from putative *Tg*Cox2a-TEV-HA parasites (clones 1 – 6). All 6 clones yielded PCR products that indicated that these clones had been successfully modified. Genomic DNA extracted from wild type (WT) parasites was used as a control.

```
TgQCR8     1 --------------------MAASRLCQYLAGRGQTGLLSLSAPRLGAPKFERKMLGSYP
PfQCR8     1 MKFNIFESKNIYNIKNKVINLKERYISFQ-----RSVVNNNNVIKKETPKFERKPLGSYP
PbQCR8     1 MVKTIFNSQAFYNIKANVADLKKRYICFQ-----RNLVRD-NIVKKETPKHEQKPLGSYP
TeQCR8     1 ---MVS----------LGASLVQRRLISGLSR--DCPNFAAKMPKRITSKHHQKPLGSYP
BbQCR8     1 --MNFF----------SALPFGQRRAFSRFAR--ASADLRASLPKRITPKHHQKPLGSYP
VbQCR8     1 -----------------------------MSLLRTRSKTVARLPTRRTSNYDQRRLGRYP
SaQCR8     1 --------------------MLLGVLWSVWSLRRAVARTSELLSLRKTSARDQLRLGSYP
AtQCR8     1 -------------------------------------------------------MGKQP
HsQCR8     1 -------------------------------------------------------MGRE-
ScQCR8     1 -------------------------------------------------------MGPP-

TgQCR8    41 VSPEFEMVWRDR-LTAHGGYIQQTISPYQLKFIYPFWHTFFARCWCKCSAYAWPWVWPG-
PfQCR8    56 IPPEAEMMWRNR-HTAYGGYIQQTISPFQQKIMYPFWHMALARWWAKFSSYLWWWIWPF-
PbQCR8    55 VPPEAEMMWRNR-HTAYGGYIQQTISPFQQKIVYPFWHMALARWWAKFSSYLWWWIWPF-
TeQCR8    46 APPEQEMLWKNR-RTIPGGYFQQSISPFQLKFLYPIIHQWSARTWAKTSQMFWWIIWPT-
BbQCR8    47 FPPEHEMLWKNR-RQVPGGYFHQAISPFQLKFCYPLIHQAYARIWAKTSQMFWWVIWPT-
VbQCR8    32 VPSEDVVWWENR-FQVPGGQWEHVVSPFQLKLWFDWAQKFPVRWYSRLSVWFWQWTYPA-
SaQCR8    41 VPPEAHIWWANR-SQVPGGEIKQTFSHLQQKIMWQWFWNAPARWYWRLSSLAWPAGVPC-
AtQCR8     6 VKL---------------KAVVYALSPFQQKIMTGLWKDLPEKIHHKVSENWISTILLVA
HsQCR8     5 --------FGN--LTRMRHVISYSLSPFEQRAYPHVFTKGIPNVLRRIRESFFRVVPQF-
ScQCR8     5 SGKTYMGWWGHMGGPKQKGITSYAVSPYAQKPLQGIFHNAVFNSFRRFKSQFLYVLIPA-

TgQCR8    99 ----LITFGLVK-KMNHDVE----------EDI--RDHYWY----
PfQCR8   114 ----AITNLILY-KMFYDAK----------KYV--KEKHWY----
PbQCR8   113 ----AITHIILW-KMFRDAK----------KYV--EQKHWY----
TeQCR8   104 ----TVMVIAFR-SIDKLNK----------LYL--KREHYY----
BbQCR8   105 ----TMAITGFL-AVEAINT----------KYV--KRIHYD----
VbQCR8    90 ----IIFYGGYF-KCCYEVD----------QSI--KAKFWY----
SaQCR8    99 ----LLMYIFVYKSCECDVE----------ASI--KAKTWW----
AtQCR8    51 PVVGTYSYAQYFKE-----Q----------EKL--EHRF------
HsQCR8    54 ----VVFYLIYTWGTE------------EFERSKRKNPAAYENDK
ScQCR8    64 ----GIYWYWWKNGNEYNEFLYSKAGREELERVNV----------
```

**Figure S2. QCR8 alignment.** Alignment of QCR8 homologs from *Toxoplasma gondii* (*Tg*QCR8; TGME49_214250)*, Plasmodium falciparum* (*Pf*QCR8; PF3D7_0306000), *P. berghei* (*Pb*QCR8; PBANKA_0404400), *Theileria equi* (*Te*QCR8; BEWA_031210), *Babesia bovis* (*Bb*QCR8; BBOV_IV004300), *Vitrella brassicaformis* (*Vb*QCR8; Vbra_14054), *Symbiodinium microadriaticum* (*Sa*QCR8; Smic7304), *Arabidopsis thaliana* (*At*QCR8; NP_196156), *Homo sapiens* (*Hs*QCR8; NP_055217) and *Saccharomyces cerevisiae* (*Sc*QCR8; NP_012369). Dark shading indicates amino acid identity in ≥70% of the sequences, and light shading indicates amino acid similarity in ≥70% of the sequences. The positions of predicted transmembrane domains in *Tg*QCR8 (TMPred prediction) and *Sc*QCR8 (TMHMM prediction) are indicated by red boxes.

```
TgQCR9     1 MHFSGVFLRTSRV-FLASE-------SSAAGSKVA--KSLPGIRFGNPWRDDYPEWIWKS
PfQCR9     1 ------------------------------------------MVFGSPFSDTYPSFIWKS
PbQCR9     1 ------------------------------------------MVFGSPFSDTYPSFIWKI
TeQCR9     1 ------------------------------------------MVFGSPFSEHYPARIWAS
BbQCR9     1 ------------------------------------------MVFGSPYSDQYPKGVWDS
SkQCR9     1 --------MLSRT-------------RLLRSALLRSKTRFSGGGFGSP-KDHEPAWIERW
VbQCR9     1 MRLSRF-HRVPRPATLTGWSSLASMYHTLLGGPLEKFYGETLGHFGSVYRQSDPHWAKAA
AtQCR9     1 ------------------------------------------------------------
ScQCR9     1 ------------------------------------------------------------
HsQCR9     1 ------------------------------------------------------------

TgQCR9    51 LRVSRKDKDMFAPFFKLLNATKLYEYCLKDNRRYCMFVMGVGLVSSWMWSEWWNSVWRRI
PfQCR9    19 LKKSRKGKDIFNPFFYALNKTKIYDHILKYNSRYWLFVVTTGCVTSYFWGVWFNNQWKRI
PbQCR9    19 LGKSRKGKDIFNPFFVALNKTRIYDHVLKYNSRYWLFVVTGGCVTSYFWGIWFNNMWKKI
TeQCR9    19 LRKSRKGKDCLSSLFHFVNSTRIYDHVLKHSKRYWIFTVFGGFASCYTISNLCDNVWKRA
BbQCR9    19 LKRSRRGMDVLSPLTSFLNSTRIYNHVLKYSKGYWLFSIAGGCMSCYAVGSVCDYVWRRV
SkQCR9    39 SANRREGQDNMDWFFRAFNATRIYEIFLKSSPRFYGFIVWSSIIGGYCWSRMWDHIWDYV
VbQCR9    60 LNSSRVGKDSMDWFFRAFNKLQLYELFLKDTPRFWLFVAVGSCLGTWYWNEAWNALWCYC
AtQCR9     1 -------MEYAARRNQKGAFEGFYKLIMRRNSVYVTFIIAGAFFGERAVDYGVHKLWERN
ScQCR9     1 -----------------MSFSSLYKTFFKRNAVFVGTIFAGAFVFQTVFDTAITSWYENH
HsQCR9     1 -------------MAAATLTSKLYSLLFRRTSTFALTIIVGVMFFERAFDQGADAIYDHI

TgQCR9   111 NKGKLYNDVPYVYPEEDE--------------------
PfQCR9    79 NKGKLYIDCPYKYPEEED--------------------
PbQCR9    79 NKGKLYIDCPYTYPEEED--------------------
TeQCR9    79 NKGKLYIDLPYQSPEED---------------------
BbQCR9    79 NRGRLYIDLPYQSPEE----------------------
SkQCR9    99 NQGKLYRHNPYVYPIPDDDE------------------
VbQCR9   120 NKGKLYRDNPYNYPEPEWDGWTPSTAPYQNFRMLTPDL
AtQCR9    54 NVGKRYEDISVLGQRPVEE-------------------
ScQCR9    44 NKGKLWKDVKARIAAGDGDDDDE---------------
HsQCR9    48 NEGKLWKHIKHKYENK----------------------
```

**Figure S3. QCR9 alignment.** Alignment of QCR9 homologs from *T. gondii* (*Tg*QCR9; TGME49_201880), *P. falciparum* (*Pf*QCR9; PF3D7_0622600), *P. berghei* (*Pb*QCR9; PBANKA_1121500), *T. equi* (*Te*QCR9; BEWA_007140), *B. bovis* (*Bb*QCR9; BBOV_III007050), *V. brassicaformis* (*Vb*QCR9; Vbra_943), *Symbiodinium kawagutii* (*Sk*QCR9; Skav217368), *Arabidopsis thaliana* (*At*QCR9; NP_190841), *Homo sapiens* (*Hs*QCR9; NP_037519) and *Saccharomyces cerevisiae* (*Sc*QCR9; NP_011699). Dark shading indicates amino acid identity in ≥70% of the sequences, and light shading indicates amino acid similarity in ≥70% of the sequences. The positions of predicted transmembrane domains in *Tg*QCR9 (TMPred prediction) and *Sc*QCR9 (TMHMM prediction) are indicated by red boxes.

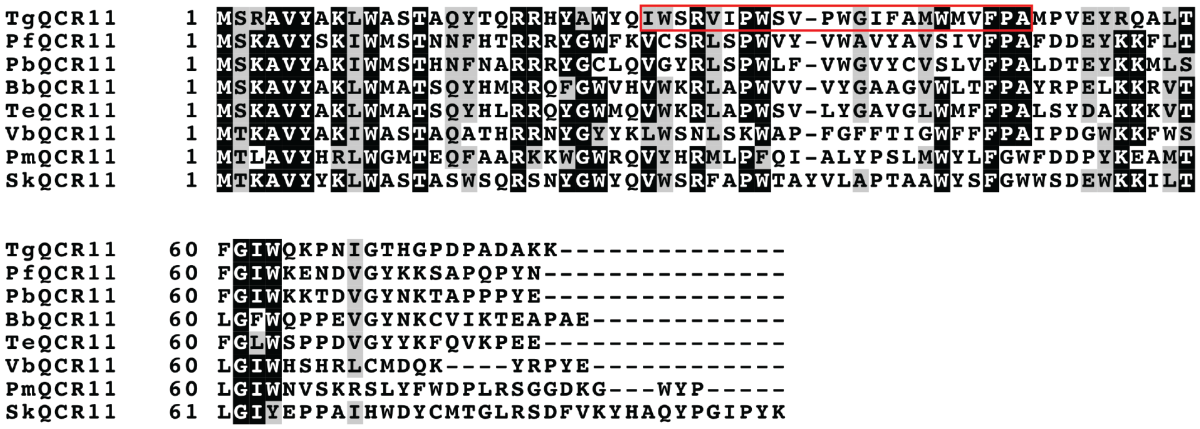


**Figure S4. QCR11 alignment.** Alignment of QCR11 homologs from *T. gondii* (*Tg*QCR11; TGME49_214250), *P. falciparum* (*Pf*QCR11; PF3D7_0722700), *P. berghei* (*Pb*QCR11; PBANKA_0620200), *B. bovis* (*Bb*QCR11; BBOV_IV004900), *T. equi* (*Te*QCR11; BEWA_032020), *V. brassicaformis* (*Vb*QCR11; Vbra_12339), *Perkinsum marinus* (*Pm*QCR11; XP_002780203), and *S. kawagutii* (*Sk*QCR11; Skav223196). Dark shading indicates amino acid identity in >70% of the sequences, and light shading indicates amino acid similarity in >70% of the sequences. The position of predicted transmembrane domains in *Tg*QCR11 (TMHMM prediction) is indicated by a red box.

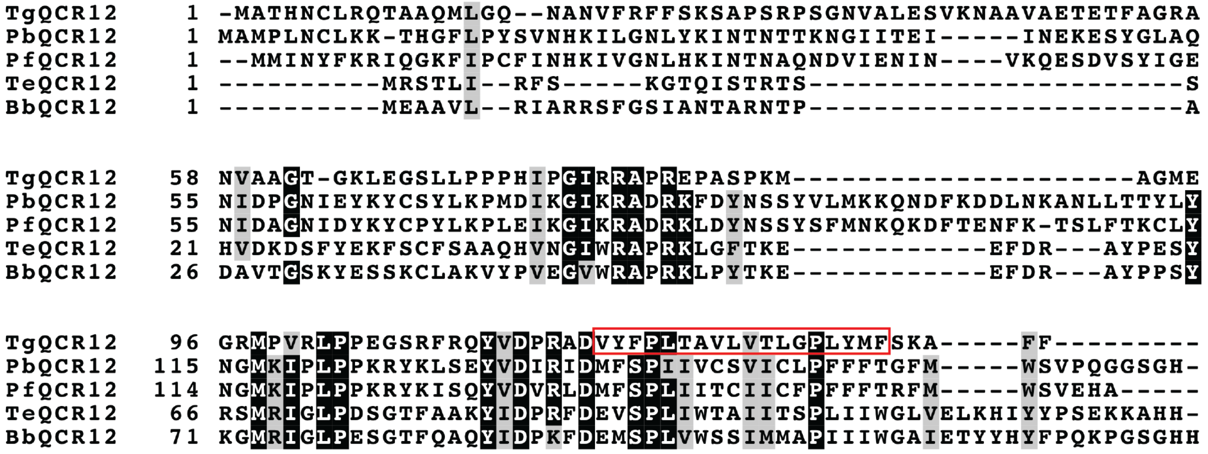


**Figure S5. QCR12 alignment.** Alignment of QCR12 homologs from *T. gondii* (*Tg*QCR12; TGME49_207170)*, P. berghei* (*Pb*QCR12; PBANKA_1341100), *P. falciparum* (*Pf*QCR12; PF3D7_1326000), *T. equi* (*Te*QCR12; BEWA_021660), and *B. bovis* (*Bb*QCR12; BBOV_III005260). Dark shading indicates amino acid identity in ≥80% of the sequences, and light shading indicates amino acid similarity in ≥80% of the sequences. The position of predicted transmembrane domains in *Tg*QCR12 (TMHMM prediction) is indicated by a red box.

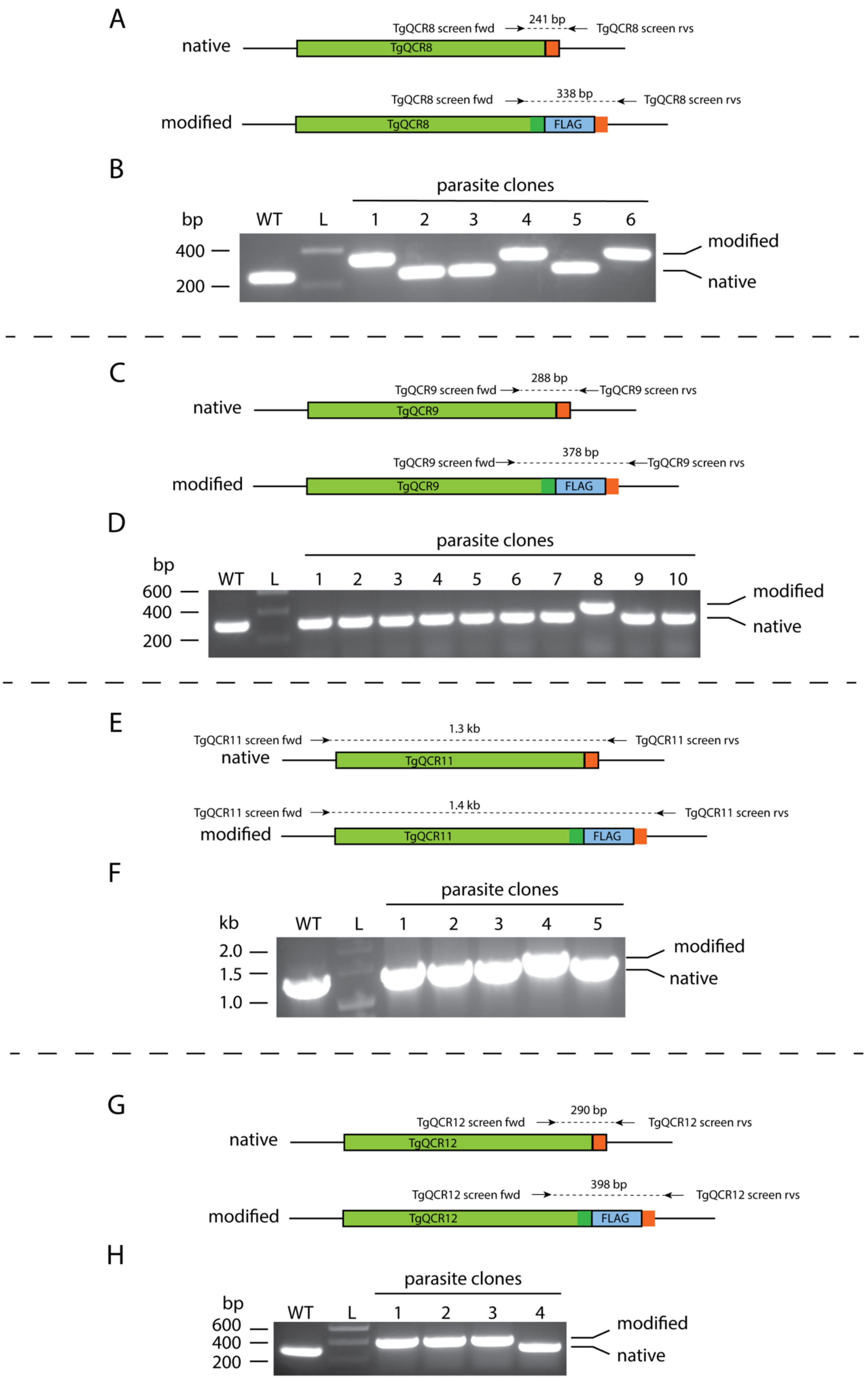
A
native
TgQCR8 screen fwd
241 bp
TgQCR8 screen rvs
TgQCR8
modified
TgQCR8 screen fwd
338 bp
TgQCR8 screen rvs
TgQCR8
FLAG
B
parasite clones
bp
WT
L
1
2
3
4
5
6
400
200
modified
native
C
native
TgQCR9 screen fwd
288 bp
TgQCR9 screen rvs
TgQCR9
modified
TgQCR9 screen fwd
378 bp
TgQCR9 screen rvs
TgQCR9
FLAG
D
parasite clones
bp
WT
L
1
2
3
4
5
6
7
8
9
10
600
400
200
modified
native
E
TgQCR11 screen fwd
1.3 kb
TgQCR11 screen rvs
native
TgQCR11
TgQCR11 screen fwd
1.4 kb
TgQCR11 screen rvs
modified
TgQCR11
FLAG
F
parasite clones
kb
WT
L
1
2
3
4
5
2.0
1.5
1.0
modified
native
G
native
TgQCR12 screen fwd
290 bp
TgQCR12 screen rvs
TgQCR12
modified
TgQCR12 screen fwd
398 bp
TgQCR12 screen rvs
TgQCR12
FLAG
H
parasite clones
bp
WT
L
1
2
3
4
600
400
200
modified
native

**Figure S6: Generating FLAG tagged *Tg*QCR8, *Tg*QCR9, *Tg*QCR11 and *Tg*QCR12 in *Tg*MPPα-HA strain parasites.** Diagrams depict the 3' replacement strategy to FLAG-tag target genes. sgRNAs were designed to target the *T. gondii* genome near the stop codon of target genes. A plasmid containing the sgRNA and GFP-tagged Cas9 endonuclease was co-transfected into *Tg*MPPα-HA *T. gondii* parasites with a PCR product encoding a FLAG epitope tag flanked by 50 bp of sequence homologous to the regions immediately up- and down-stream of the stop codon. Genomic DNA extracted from wild type (WT) parasites was used as a control in PCRs. **(A)** Forward and reverse primers were used to screen parasite clones for integration of the FLAG tag at the *Tg*QCR8 locus, yielding a 241 bp product in the native locus and a 338 bp product in the modified locus. **(B)** PCR screening using genomic DNA extracted from putative *Tg*QCR8-FLAG parasites (clones 1 – 6). Clones 1, 4 and 6 yielded PCR products that indicated that these clones had been successfully modified. **(C)** Forward and reverse primers were used to screen parasite clones for integration of the FLAG tag at the *Tg*QCR9 locus, yielding a 288 bp product in the native locus and a 378 bp product in the modified locus. **(D)** PCR screening using genomic DNA extracted from putative *Tg*QCR9-FLAG parasites (clones 1 – 10). Clone 8 yielded a PCR product that indicated it had been successfully modified. **(E)** Forward and reverse primers were used to screen parasite clones for integration of the FLAG tag at the *Tg*QCR11 locus, yielding a 1.3 kb product in the native locus and a 1.4 kb product in the modified locus. **(F)** PCR screening using genomic DNA extracted from putative *Tg*QCR11-FLAG parasites (clones 1 – 5). Clone 4 yielded a PCR product that indicated it had been successfully modified. **(G)** Forward and reverse primers were used to screen parasite clones for integration of the FLAG tag at the *Tg*QCR12 locus, yielding a 290 bp product in the native locus and a 398 bp product in the modified locus. **(H)** PCR screening using genomic DNA extracted from putative *Tg*QCR12-FLAG parasites (clones 1 – 4). Clones 1-3 yielded PCR products that indicated that these clones had been successfully modified.

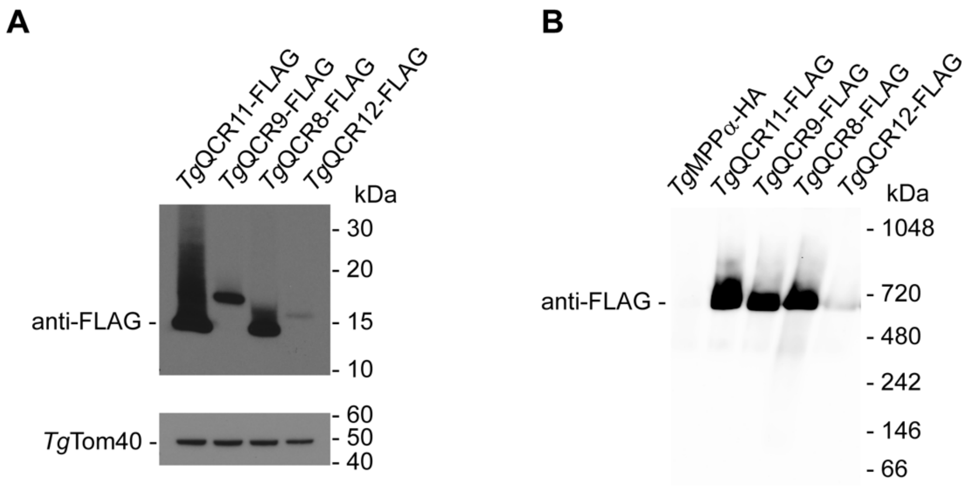


**Figure S7: Candidate Complex III subunits are expressed at different levels in *T. gondii* parasites.** **(A)** Western blot of proteins extracted from *Tg*MPPα-HA/*Tg*QCR11-FLAG, *Tg*MPPα-HA/*Tg*QCR9-FLAG, *Tg*MPPα-HA/*Tg*QCR8-FLAG and *Tg*MPPα-HA/*Tg*QCR12-FLAG parasites, separated by SDS-PAGE, and detected with anti-FLAG and anti-Tom40 (loading control) antibodies. **(B)** Western blot of proteins extracted from *Tg*MPPα-TEV-HA, *Tg*MPPα-HA/*Tg*QCR11-FLAG, *Tg*MPPα-HA/*Tg*QCR9-FLAG, *Tg*MPPα-HA/*Tg*QCR8-FLAG and *Tg*MPPα-HA/*Tg*QCR12-FLAG parasites, separated by BN-PAGE, and detected with anti-FLAG antibodies.

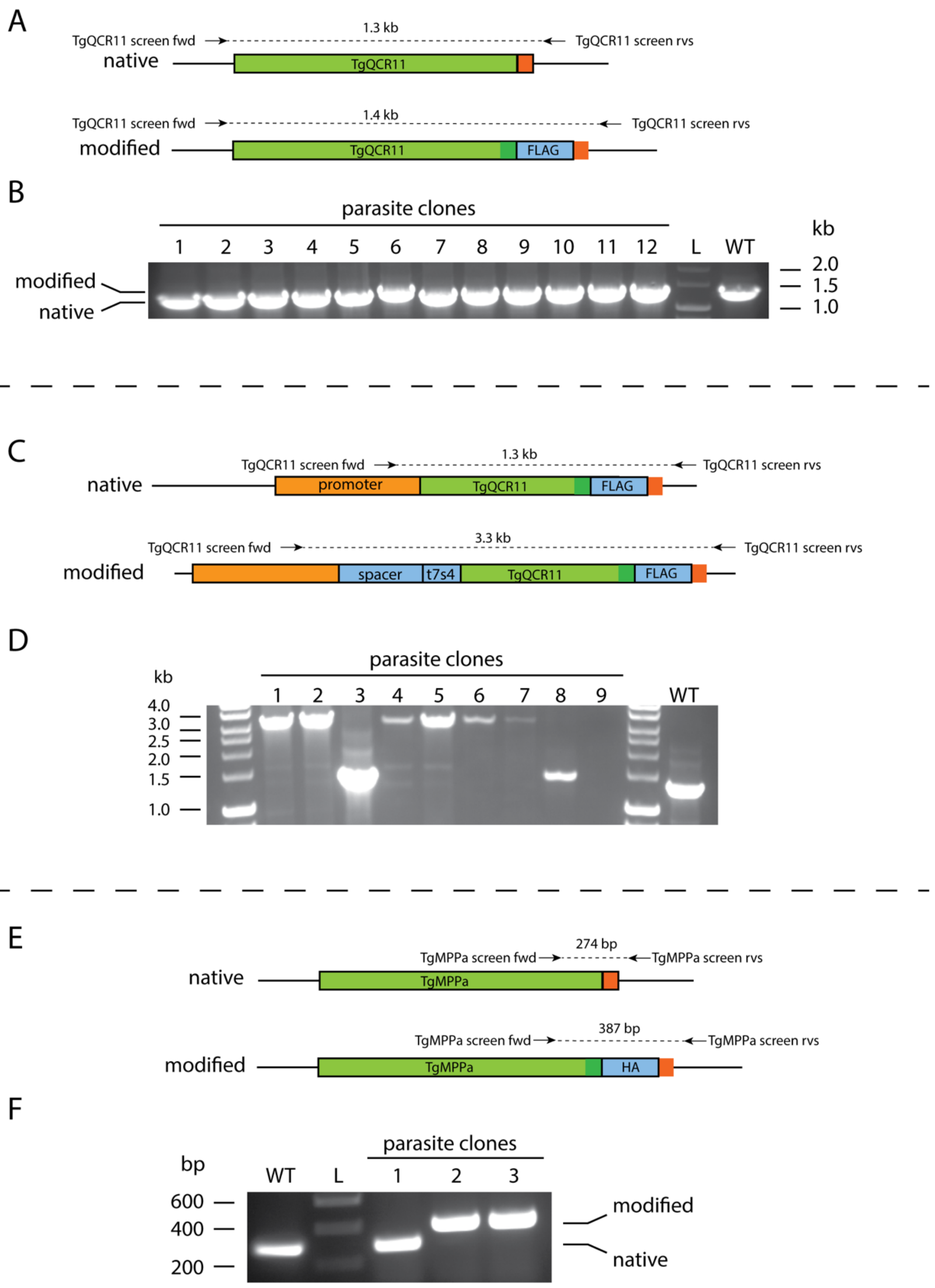

A
TgQCR11 screen fwd
1.3 kb
TgQCR11 screen rvs
native
TgQCR11
TgQCR11 screen fwd
1.4 kb
TgQCR11 screen rvs
modified
TgQCR11
FLAG
B
parasite clones
1 2 3 4 5 6 7 8 9 10 11 12 L WT
kb
2.0
1.5
1.0
modified
native
C
TgQCR11 screen fwd
1.3 kb
TgQCR11 screen rvs
native
promoter
TgQCR11
FLAG
TgQCR11 screen fwd
3.3 kb
TgQCR11 screen rvs
modified
spacer
t7s4
TgQCR11
FLAG
D
parasite clones
kb
1 2 3 4 5 6 7 8 9 WT
4.0
3.0
2.5
2.0
1.5
1.0
E
274 bp
TgMPPa screen fwd
TgMPPa screen rvs
native
TgMPPa
387 bp
TgMPPa screen fwd
TgMPPa screen rvs
modified
TgMPPa
HA
F
parasite clones
bp
WT L 1 2 3
600
400
200
modified
native

**Figure S8: Generating an ATc regulated, FLAG-tagged *Tg*QCR11 and r*Tg*QCR11-FLAG/*Tg*MPPα-HA parasite strains. (A)** Diagram depicting the 3' replacement strategy to FLAG-tag *Tg*QCR11. A sgRNA was designed to target the *T. gondii* genome near the stop codon of *Tg*QCR11. A plasmid containing the sgRNA and GFP-tagged Cas9 endonuclease was co-transfected into *T. gondii* parasites with a PCR product encoding a FLAG epitope tag flanked by 50 bp of sequence homologous to the regions immediately up- and down-stream of the *Tg*QCR11 stop codon. Forward and reverse primers were used to screen parasite clones for integration of the FLAG tag at the *Tg*QCR11 locus, yielding a 1.3 kb product in the native locus and a 1.4 kb product in the modified locus. **(B)** PCR screening using genomic DNA extracted from putative *Tg*QCR11-FLAG parasites (clones 1 – 12). Clone 6 yielded a PCR product that indicated it had been successfully modified. Genomic DNA extracted from wild type (WT) parasites was used as a control. **(C)** Diagram depicting the promoter replacement strategy to generate ATc-regulated *Tg*QCR11. A sgRNA was designed to target the *T. gondii* genome near the start codon of *Tg*QCR11. A plasmid containing the sgRNA and GFP-tagged Cas9 endonuclease was co-transfected into *T. gondii* parasites with a PCR product encoding the ATc regulated 't7s4' promoter, which contains 7 copies of the Tet operon and a Sag4 minimal promoter, flanked by 50 bp of sequence homologous to the regions immediately up- and down-stream of the *Tg*QCR11 start codon. The PCR product also contain a 'spacer' region that separates the regulatable promoter from the native promoter of the *Tg*QCR11 gene to enable sufficient regulation. Forward and reverse primers were used to screen parasite clones for successful integration of the regulatable promoter at the *Tg*QCR11 locus, yielding a 1.3 kb product in the native locus and a 3.3 kb product in the modified locus. **(D)** PCR screening using genomic DNA extracted from putative r*Tg*QCR11-FLAG parasites (clones 1 – 9). Clones 1, 2, 4-7 yielded PCR products that indicated that these clones had been successfully modified. Genomic DNA extracted from wild type (WT) parasites was used as a control. **(E)** Diagram depicting the 3' replacement strategy to HA-tag *Tg*MPPα. A sgRNA was designed to target the *T. gondii* genome near the stop codon of *Tg*MPPα. A plasmid containing the sgRNA and GFP-tagged Cas9 endonuclease was co-transfected into r*Tg*QCR11-FLAG *T. gondii* parasites with a PCR product encoding a HA epitope tag flanked by 50 bp of sequence homologous to the regions immediately up- and down-stream of the *Tg*MPPα stop codon. Forward and reverse primers were used to screen parasite clones for integration of the HA tag at the *Tg*MPPα locus, yielding a 274 bp product in the native locus and a 387 bp product in the modified locus. **(F)** PCR screening using genomic DNA extracted from putative r*Tg*QCR11/*Tg*MPPα-HA parasites (clones 1 – 3). Clones 2 and 3 yielded PCR products that indicated that these clones had been successfully modified. Genomic DNA extracted from wild type (WT) parasites was used as a control.

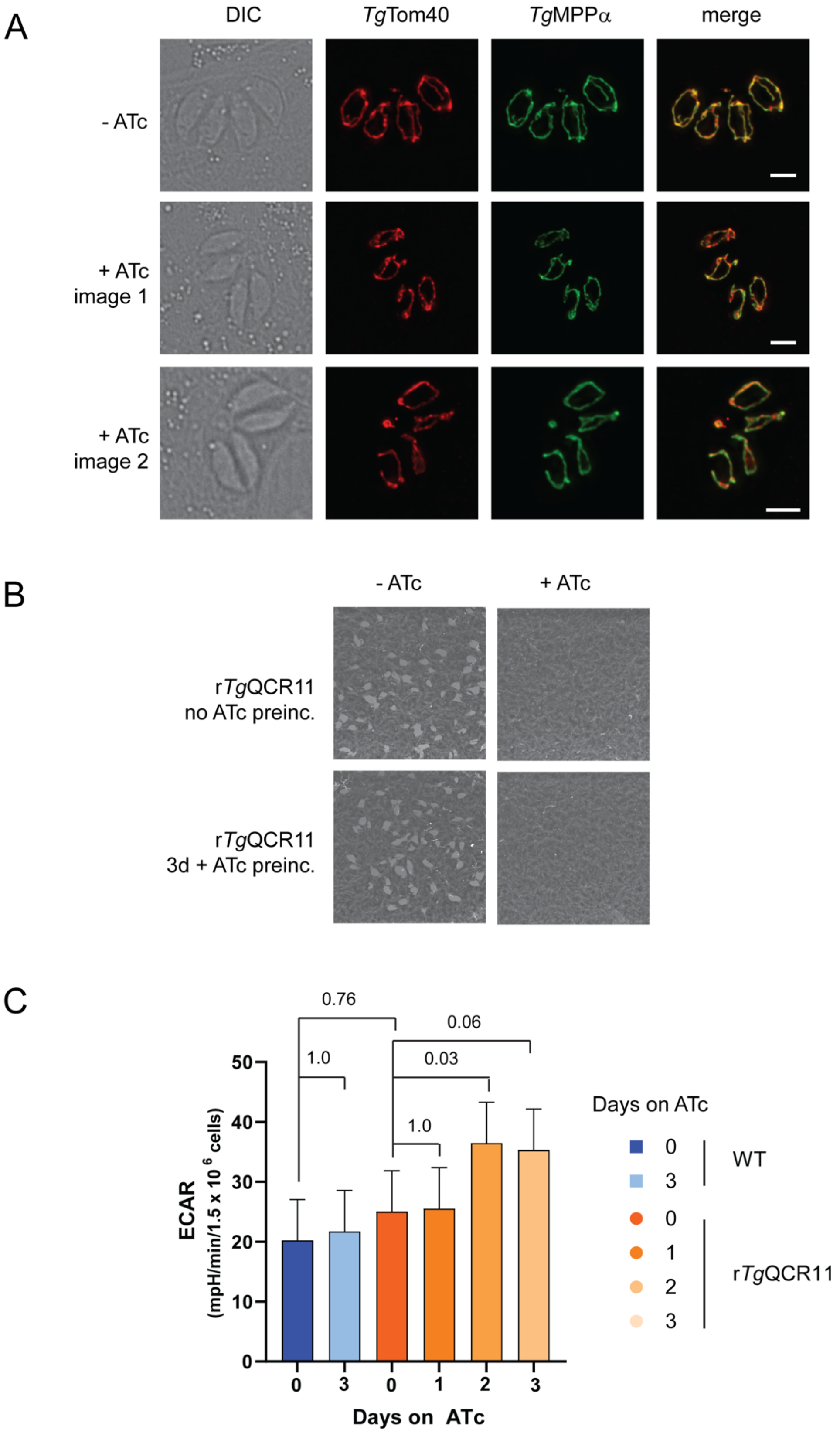
A
DIC
TgTom40
TgMPPα
merge
- ATc
+ ATc
image 1
+ ATc
image 2
B
- ATc
+ ATc
rTgQCR11
no ATc preinc.
rTgQCR11
3d + ATc preinc.
C
0.76
0.06
1.0
0.03
1.0
ECAR
(mpH/min/1.5 x 10 6 cells)
50
40
30
20
10
0
0
3
0
1
2
3
Days on ATc
Days on ATc
0
3
WT
0
1
2
3
rTgQCR11

**Figure S9: Defects in mOCR observed upon knockdown of *Tg*QCR11 are not caused by general defects in mitochondrial morphology, parasite viability or parasite metabolism.** **(A)** Immunofluorescence assays assessing mitochondrial morphology in r*Tg*QCR11-FLAG/*Tg*MPPα-HA parasites grown in the absence of ATc (top) or in the presence of ATc for 3 days (middle and bottom). The outer mitochondrial membrane was labelled using antibodies against *Tg*Tom40 (red), and the inner mitochondrial membrane was labelled using anti-HA antibodies to detect *Tg*MPPα-HA. Images are representative of 100 four-cell vacuoles examined in 2 independent experiments; scale bar represents 2 μm. **(B)** Plaque assays of r*Tg*QCR11-FLAG/*Tg*MPPα-HA parasites grown for 8 days in the absence (left) or presence (right) of ATc. r*Tg*QCR11 parasites were either grown in the absence of ATc (no ATc preinc; top) or pre-incubated in ATc for 3 days (3d + ATc preinc; bottom) before commencing the experiment. Plaque assays are from a single experiment, representative of 3 independent experiments. **(C)** Basal extracellular acidification rate (ECAR) of WT parasites grown in the absence of ATc or in the presence of ATc for 3 days (blue), and r*Tg*QCR11-FLAG/*Tg*MPPα-HA parasites grown in the absence of ATc or in the presence of ATc for 1-3 days (orange). A linear mixed-effects model was fitted to the data and values depict the least squares mean ± 95% CI of three independent experiments. ANOVA followed by Tukey's multiple pairwise comparisons test was performed, with relevant $p$ values shown.

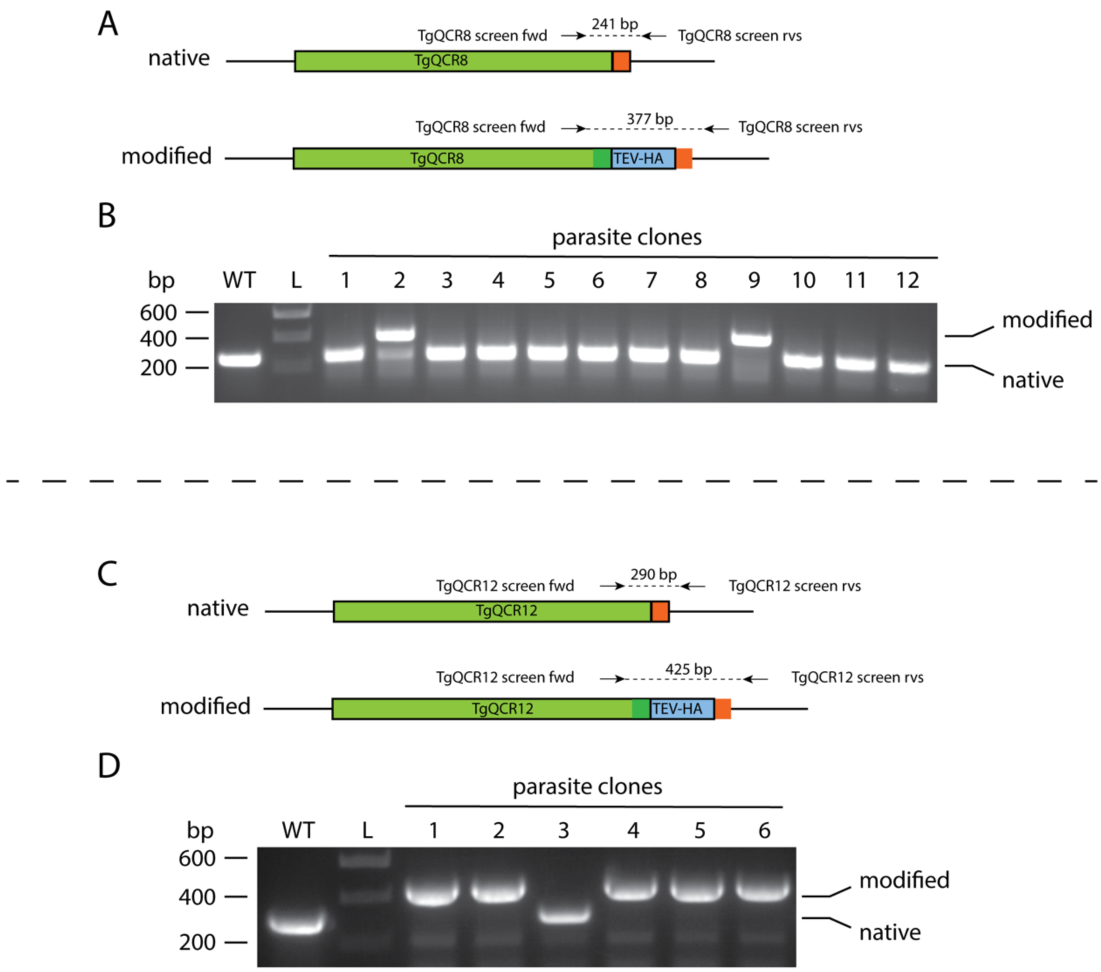
A
native
TgQCR8
TgQCR8 screen fwd
241 bp
TgQCR8 screen rvs
modified
TgQCR8
TEV-HA
TgQCR8 screen fwd
377 bp
TgQCR8 screen rvs
B
parasite clones
bp
WT
L
1
2
3
4
5
6
7
8
9
10
11
12
600
400
200
modified
native
C
native
TgQCR12
TgQCR12 screen fwd
290 bp
TgQCR12 screen rvs
modified
TgQCR12
TEV-HA
TgQCR12 screen fwd
425 bp
TgQCR12 screen rvs
D
parasite clones
bp
WT
L
1
2
3
4
5
6
600
400
200
modified
native

**Figure S10: Generating TEV-HA tagged *Tg*QCR8 and *Tg*QCR12 in r*Tg*QCR11-FLAG parasites.** Diagrams depict the 3’ replacement strategy to TEV-HA-tag target genes. sgRNAs were designed to target the *T. gondii* genome near the stop codon of target genes. A plasmid containing the sgRNA and GFP-tagged Cas9 endonuclease was co-transfected into r*Tg*QCR11-FLAG *T. gondii* parasites with a PCR product encoding a TEV-HA epitope tag flanked by 50 bp of sequence homologous to the regions immediately up- and down-stream of the stop codon. Genomic DNA extracted from wild type (WT) parasites was used as a control in PCRs. **(A)** Forward and reverse primers were used to screen parasite clones for integration of the TEV-HA tag at the *Tg*QCR8 locus, yielding a 241 bp product in the native locus and a 377 bp product in the modified locus. **(B)** PCR screening using genomic DNA extracted from putative r*Tg*QCR11-FLAG/*Tg*QCR8-TEV-HA parasites (clones 1 – 12). Clone 9 yielded PCR products that indicated it had been successfully modified. **(C)** Forward and reverse primers were used to screen parasite clones for integration of the TEV-HA tag at the *Tg*QCR12 locus, yielding a 290 bp product in the native locus and a 425 bp product in the modified locus. **(D)** PCR screening using genomic DNA extracted from putative r*Tg*QCR11-FLAG/*Tg*QCR12-TEV-HA parasites (clones 1 – 6). Clones 1-2 and 4-6 yielded PCR products that indicated that they had been successfully modified.

**Table S1: Extended data from the mass spectrometry-based proteomic analysis of the *Tg*MPPα complex.** Tab 1: List of proteins identified in the mass spectrometry-based proteomic analysis of proteins purified from the *Tg*MPPα-TEV-HA and *Tg*Cox2a-TEV-HA immunoprecipitations. Included are the ToxoDB (www.toxodb.org) accession number, the name assigned to the protein in this manuscript if applicable, the total precursor intensity (area) for each biological replicate, the average total precursor intensity (area) of the three replicates, the fold change (average MPPα/average Cox2a), and the predicted mass of the identified protein. Proteins are highlighted based on whether they were highly enriched in TgMPPα-TEV-HA (orange) or TgCox2a-TEV-HA (blue) pulldown. Tab 2: A list of the log fold change (logFC) and *p* values calculated for each protein identified in all replicates of the *Tg*MPPα-TEV-HA and *Tg*Cox2a-TEV-HA immunoprecipitations following EdgeR analysis.

**Table S2.** Sequences of oligonucleotides (primers) and gBlocks used in this study.

| Oligonucleotide name | Oligonucleotide sequence (5' to 3') |
|---|---|
| MPPα 3' rep CRISPR fwd | GCTTCACTTGCCGACGCCCGGTTTTAGAGCTAGAAATAGCAAG |
| Universal CRISPR rvs | ACTTGACATCCCCATTTAC |
| MPPα tag fwd | CGCACTACGAGGAGGTACGCGCTGCTCTCCGAGCAGCGGGCGTCGGCAAG GGTGGAGGTAGCGGTGGTGGAAG |
| MPPα tag rvs | ATGCAGCTTTCTTCGTTTCCGAGACCTTTCCAATTCTCTCTGCGCCCTGCGC TTCTGTGGGCGGTTATCAGG |
| Cox2a tag fwd | GACAGTGGTACTGGATCTACGAAGTCGAGTCGCCTGTTGACGACGAaGAGG GTGGAGGTAGCGGTGGTGGAAG |
| Cox2a tag rvs | CTGCCCATTCAACGCTCGGACAGCCGTCCTTTAGGAAACGCATAGGAAGCG CTTCTGTGGGCGGTTATCAGG |
| MPPα screen fwd | TTTCTTTTTCGCTGTCCGATA |
| MPPα screen rvs | GTAGACACGTTCCTTCCTCTCG |
| Cox2a screen fwd | CTCTTGTACATGCTCGACGAAG |
| Cox2a screen rvs | AACGACTGTGATTCCAAAACCT |
| QCR8 3' rep CRISPR fwd | TCGCAGGTATTAAGCGTCGTGTTTTAGAGCTAGAAATAGCAAG |
| QCR9 3' rep CRISPR fwd | CCGGAGGAGGATGAGTAAACGTTTTAGAGCTAGAAATAGCAAG |
| QCR11 3' rep CRISPR fwd | CACTGTCTATTTCTTTGCGTGTTTTAGAGCTAGAAATAGCAAG |
| QCR12 3' rep CRISPR fwd | AAGCTGGTTCTACAGTGCCGGTTTTAGAGCTAGAAATAGCAAG |
| QCR8 tag fwd | GAACTGGAAGAGAAGTGCGTCCTTTTTCTGGGTGTTTCGCTCGCAGGTATG GTGGAGGTAGCGGTGGTGGAAG |
| QCR8 tag rvs | CAGGGTTTCTCTCCCGCAAAGAGGCGAATCTGGACCGGCAAACATCGTTGG CTTCTGTGGGCGGTTATCAGG |
| QCR9 tag fwd | AACAGAAACTCTACAATGATGTCCCGTACGTCTATCCGGAGGAGGATGAG GGTGGAGGTAGCGGTGGTGGAAG |
| QCR9 tag rvs | GCCACGGGTGAGTGGAACCTTGCAGCTTCAGATTCGTGTTTGCATAGCAGG CTTCTGTGGGCGGTTATCAGG |
| QCR11 tag fwd | AAAAACCGAACATTGGCACTCACGGACCGGATCCTGCtGACGCAAAGAAA GGTGGAGGTAGCGGTGGTGGAAG |
| QCR11 tag rvs | GAGTTTGGCGACCGTGTTGTTGAGCCGTACTTCCCAATGAAATGCTCTCGG CTTCTGTGGGCGGTTATCAGG |
| QCR12 tag fwd | CTGTTTTGGTGACGCTGGGGCCTCTCTACATGTTCTCCAAAGCCTTCTTCGG TGGAGGTAGCGGTGGTGGAAG |
| QCR12 tag rvs | TCTCGCGGCTTTCTTGGAGTTCGCGCCCCAGAGGAGAGACTCGCACCACGG CTTCTGTGGGCGGTTATCAGG |
| QCR8 screen fwd | GTCTTCAGGGTCTTCTGTTGCT |
| QCR8 screen rvs | CTTCCGTTTTACGAGCTCAAGT |
| QCR9 screen fwd | CGTTTTCACACACACTACCCAT |
| QCR9 screen rvs | TGACTTGTGTTGCAGAGTAGGC |
| QCR11 screen fwd | TTTTTATCTATTCTGGGCCTGC |
| QCR11 screen rvs | CCCATACCTCACTGGTTTCTGT |
| QCR12 screen fwd | CGGACGTTTACTTTCCTCTCAC |
| QCR12 screen rvs | TGAAACAGTGTCCAGAGACGAC |
| QCR11 5' CRISPR fwd | GGACATTCTGGCTCCGGCAGGTTTTAGAGCTAGAAATAGCAAG |
| QCR11 pro rep fwd | CGAGTTTTTTTCTGCCGAACAGGCGGTTATTCTCAAGGTAATTTCTCCCAGG TTGCAGGCTCCTTCTTCGG |
| QCR11 pro rep rvs | GTGTATTGAGCCGTGCTGGCCCAGAGCTTCGCGTAGACCGCGCGGGACATtt tGGTTGAAGACAGACGAAAGCAGTTG |
| QCR11 comp fwd | CTAGAGATCTAAAATGTCCCGAGCGGTCTACGC |
| Universal Ty1 rvs | CTAGCCCGGGGCTTCTGTGGGCGGTTATCAGG |

| | |
|---|---|
| TEV-HA gBlock | GGTGGAGGTAGCGGTGGTGGAAGTGAAAATCTGTACTTCCAGGGAGGTACCTACCCGTACGACGTCCCGGACTACGCTGGCTATCCCTATGATGTGCCCGATTATGCGTATCCTTACGATGTTCCAGATTATGCCTGATAACCGCCCACAGAAGC |
| FLAG gBlock | GTGGAGGTAGCGGTGGTGGAAGTGACTACAAAGACCATGACGGTGATTATAAAGATCATGACATCGATTACAAGGATGACGATGACAAGTAGTCCTGATAACCGCCCACAGAAGC |
| HA gBlock | GGTGGAGGTAGCGGTGGTGGAAGTTACCCGTACGACGTCCCGGACTACGCTGGCTATCCCTATGATGTGCCCGATTATGCGTATCCTTACGATGTTCCAGATTATGCCTGATAACCGCCCACAGAAGC |
| QCR11-Ty1 gBlock | ATGTCCCGAGCGGTCTACGCGAAGCTCTGGGCCAGCACGGCTCAATACACACAACGCAGACATTATGCGTGGTACCAAATCTGGTCGCGCGTGATTCCCTGGTCCGTGCCTTGGGGCATCTTCGCTATGTGGATGGTGTTCCCCGCCATGCCAGTTGAGTATCGTCAGGCGCTGACTTTCGGCATTTGGCAAAAACCGAACATTGGCACTCACGGACCGGATCCTGCCGACGCAAAGAAAGGTGGAGGTAGCGGTGGTGGAAGTGAGGTGCATACCAATCAAGACCCTTTGGATGAAGTCCATACCAATCAAGATCCTTTGGACGAGGTCCATACGAACCAGGACCCCTTGGACGGGGCCTGATAACCGCCCACAGAAGC |
